# Genetically Encoded CRISPR components Yield Efficient Gene Editing in the Invasive Pest, *Drosophila suzukii*

**DOI:** 10.1101/2021.03.15.435483

**Authors:** Nikolay P. Kandul, Esther J. Belikoff, Junru Liu, Anna Buchman, Fang Li, Akihiko Yamamoto, Ting Yang, Isaiah Shriner, Maxwell J. Scott, Omar S. Akbari

## Abstract

Originally from Asia, *Drosophila suzukii* (Matsumura, 1931, Diptera: *Drosophilidae*) is presently a global pest of economically important soft-skinned fruits. Also commonly known as spotted wing *Drosophila* (SWD), it is largely controlled through repeated applications of broad-spectrum insecticides. There is a pressing need for a better understanding of SWD biology and for developing alternative environmentally-friendly methods of control. The RNA-guided Cas9 nuclease has revolutionized functional genomics and is an integral component of several recently developed genetic strategies for population control of insects. Here we have developed transgenic strains that encode three different terminators and four different promoters to express Cas9 in both the soma and/or germline of SWD. The Cas9 lines were evaluated through genetic crossing to transgenic lines that encode single guide RNAs targeting the conserved X-linked *yellow* body and *white* eye genes. We find that several Cas9/gRNA lines display very high editing capacity. Going forward, these tools will be instrumental for evaluating gene function in SWD and may provide tools useful for the development of new genetic strategies for control of this invasive species.

## Introduction

*Drosophila suzukii*, commonly known as Spotted Wing Drosophila (SWD), is a significant crop pest of many soft-skinned fruits that has recently invaded much of the world (1–3). Unlike other fly species that infest overripe, or rotting fruits, SWD targets ripening fruits (1, 3, 4). SWD females (♀) use a serrated ovipositor to pierce the fruit skin, and deposit progeny inside to consume the fruit (1). The external wounds generated by oviposition alone leave the fruit vulnerable to secondary infections caused by pathogens including bacteria, yeasts, and fungi. The short generation time of SWD contributes to its rapid infestation, resulting in significant revenue losses (5–7). Endemic to East Asia, SWD has become established around much of Europe, North America, and South America since 2008 (6–9), with modeling predicting even further spread (10). Given the invasiveness of SWD, and the significant crop damages, there exists a pressing need to generate molecular tools that can be used for both gaining a better understanding of SWD biology and to innovate alternative environmentally-friendly control methodologies.

SWD is largely controlled through the use of insecticides (11–14). However, insecticide applications can provide limited protection, indiscriminately affect beneficial species (15), and insecticide resistance has emerged (16). Environmentally-friendly species-specific methods of insect pest control, such as the sterile insect technique (SIT), and incompatible insect technique (IIT), are being developed for SWD. In SIT applications, flies are mass reared and exposed to ionizing radiation for sterilization. Then, excess sterilized males (♂’s) are repeatedly released to mate with wild ♀’s, which in turn lay unfertilized eggs (17). While radiation conditions for SWD have been identified that produce sterile ♂’s and ♀’s (18, 19), further testing is necessary to determine if these ♂’s are competitive and can suppress populations. In IIT applications, flies are instead infected with the endosymbiont *Wolbachia*, which has been shown to induce cytoplasmic incompatibility (CI) when infected ♂’s mate with non-infected ♀’s. Here, the paternal sperm is no longer recognized by the egg, leaving the egg unfertilized (20, 21). While IIT can be a promising application, accidental release of infected ♀’s can lead to unintended replacement of a wild population with an infected/resistant one, as CI does not occur when infected ♀’s mate with either infected or non-infected ♂’s (22). Further, both naturally occurring and trans-infected *Wolbachia* strains do not provide 100% CI in SWD, which is required for an effective IIT (23). The combination of SIT and IIT was recently assessed to improve the sterility of *Wolbachia* infected SWD. However, the mating competitiveness of released SWD ♂’s was not assessed (24). Thus, while progress has been made, it remains to be determined if SIT and/or IIT can provide an alternative economical means for control of SWD.

Over the past several years there has been significant progress in developing molecular genetic tools that can be used for both gaining a better understanding of SWD biology and to innovate alternative environmentally-friendly control methods. For instance, transgenesis has been achieved by several groups (25–28), a high quality reference genome has been assembled (29), and the versatile RNA-guided CRISPR/Cas9 nuclease (clustered regularly interspaced short palindromic repeats/CRISPR-associated sequence 9) has recently been used for gene editing (30, 31). For gene editing in SWD, site-specific mutations were initially made through microinjection of embryos with Cas9/gRNA plasmid DNAs (31). The efficiency of mutagenesis was later improved upon by microinjection with recombinant Cas9 protein and synthetic gRNA rather than plasmid DNAs (30). This editing efficiency could be further improved by generating transgenic strains that encode Cas9/gRNA, as done in *D. melanogaster* (32–35). Further, development of effective Cas9/gRNA strains may also optimize site-specific transgenesis methods (36, 37), and prove useful for generating a variety of genetic control systems in SWD such as precision-guided SIT (pgSIT) (38, 39), or possibly even homing based gene drives (40, 41) for population suppression.

Given the many advantages of generating transgenic lines encoding Cas9/gRNAs, here we report the development and evaluation of such strains in SWD. In total we generated 8 homozygous Cas9 expressing lines that were driven by four separate promoters (*nos, vasa, BicC*, and *ubiq*) using three different terminators to provide robust expression in both the soma and/or germline. To evaluate the efficacy of these strains, we also generated strains encoding single guide RNAs that target two conserved X-linked recessive genes known to produce visible phenotypes when disrupted including *yellow* body and *white* eye genes. By crossing the Cas9 and gRNA strains together, we find that several strains display remarkably high rates of editing capacity. Going forward, these tools will be invaluable for characterization of gene function and may also prove useful for engineering novel control strategies for this invasive crop pest.

## Results

### Development of Cas9 and gRNA encoding strains

The aim of the study was to develop a versatile toolbox for gene editing and genome engineering in SWD by generating transgenic strains encoding CRISPR components (i.e. Cas9 and gRNAs). To robustly express and import Cas9 into nuclei, we used *Streptococcus pyogenes Cas9* (*Cas9*) with a nuclear localization sequence (NLS) on either the C-terminal end only (*Cas9-NLS (32))*, or on both terminals (*NLS-Cas9-NLS (42)*) (**Fig. 1A**). To drive expression of Cas9, we used *D. melanogaster* promoters expressed in either early germ cells, *vasa* (*vas*) (43) or *nanos* (*nos*) (44), or in late germ cells, *Bicaudal C* (*BicC*) (45), or in both germ and somatic cells, *polyubiquitin 63E* (*ubiq) (46)*. Using each promoter, we built four *piggyBac* constructs that express *NLS-Cas9-NLS* terminated by a p10 3′-UTR derived from the *Autographa californica* nucleopolyhedrovirus (AcNPV) (47, 48) for strong translation of Cas9, and a red (*Opie2-dsRed*) transgenesis marker (referred as *Cas9*.*R*). We also built two alternative *piggyBac* constructs that contain either the *vas* or *nos* promoters driving expression of the *Cas9-NLS* terminated with *vas* or *nos* 3’UTR’s from *D. melanogaster*, with a green (*ubiq-ZsGreen*) transgenesis marker (referred as *Cas9*.*G*) (**Fig. 1A**). In total, six Cas9 constructs were engineered that were used to generate 8 homozygous transgenic strains (at least one homozygous transgenic strain per construct), two of which were X-linked (**Fig. 1B, Table 1**).

**Table 1.** SWD transgenic strains generated in the study

| Strain | Construct ID | Strain marker | Sequence reference | Strain availability | Additional information |
| --- | --- | --- | --- | --- | --- |
| **vasCas9.R** | vasCas9.R (874Z) | Opie2-dsRed | AddGene #112687 | O. Akbari at UCSD | Cas9 expressed in somatic and germ cells |
| **BicC.Cas9.R** | BicC.Cas9.R (874R) | Opie2-dsRed | AddGene #168295 | O. Akbari at UCSD | Cas9 expressed in late germ cells |
| **ubiqCas9.R** | ubiqCas9.R (874W) | Opie2-dsRed | AddGene #112686 | O. Akbari at UCSD | X-linked Cas9 expressed in somatic and germ cells |
| **nosCas9.R** | nosCas9.R (874Z1) | Opie2-dsRed | AddGene #112685 | O. Akbari at UCSD | Cas9 expressed in somatic and germ cells |
| **vasCas9.G#4** | vasCas9.G | ubiq-ZsGreen | Addgene# 169012 | M. Scott at NCSU | Cas9 expressed in germ cells |
| **vasCas9.G#6** | vasCas9.G | ubiq-ZsGreen | Addgene# 169012 | M. Scott at NCSU | Cas9 expressed in germ cells |
| **nosCas9.G#14** | nosCas9.G | ubiq-ZsGreen | Addgene# 169011 | M. Scott at NCSU | Cas9 expressed in germ cells |
| **nosCas9.G#36** | nosCas9.G | ubiq-ZsGreen | Addgene# 169011 | M. Scott at NCSU | X-linked Cas9 expressed in germ cells |
| **gRNA<sup>y</sup>** | gRNA<sup>y</sup> (982G.2) | Opie2-eGFP, 3xP3-eGFP | AddGene #168294 | O. Akbari at UCSD | gRNA targets *yellow* exon #2 |
| **gRNA<sup>w</sup>** | gRNA<sup>w</sup> | ubiq-dsRed | Addgene# 169010 | M. Scott at NCSU | gRNA targets *white* exon #2 |

**Figure 1.**
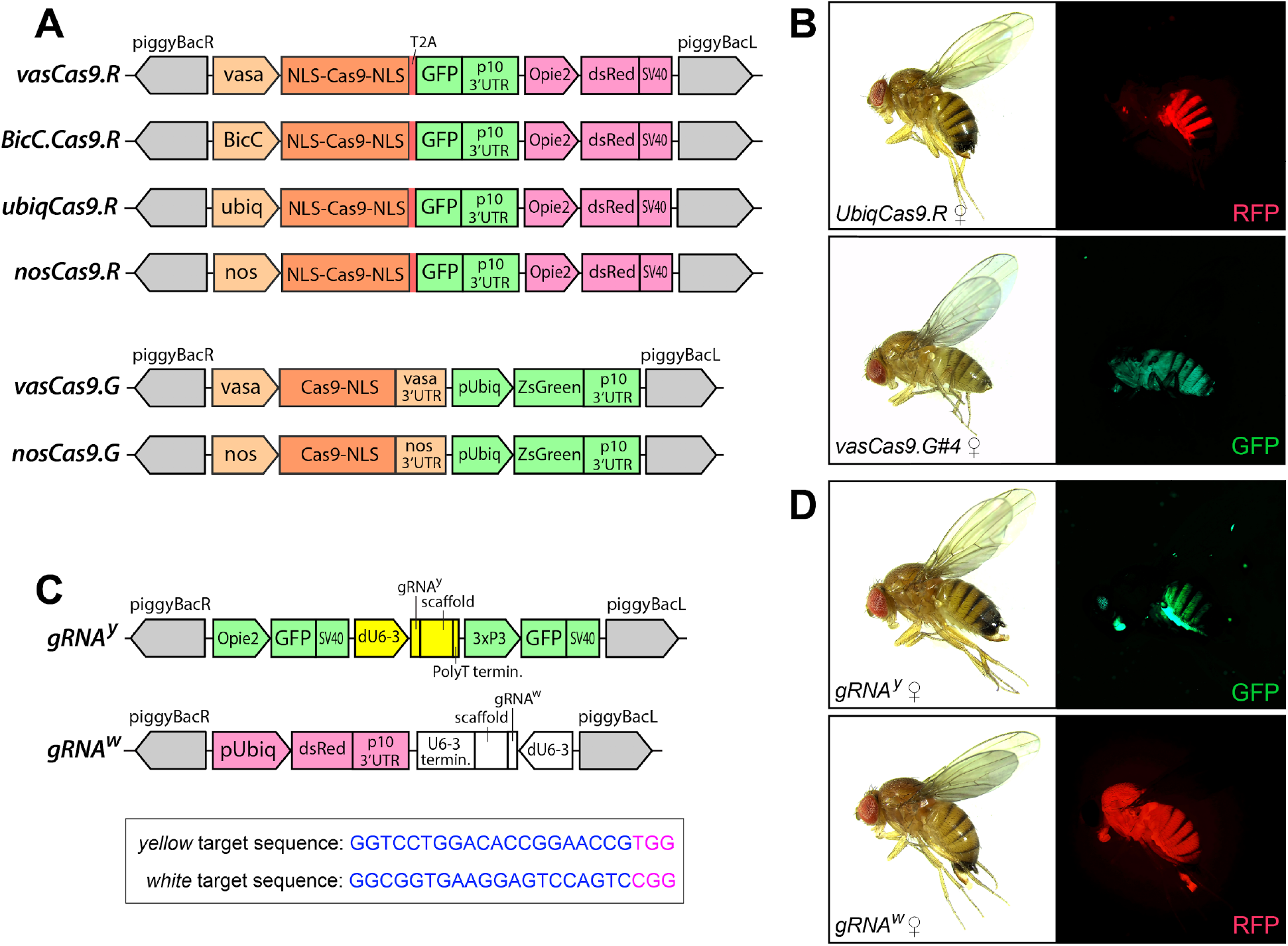
Schematic maps of genetic constructs and images of transgenic SWD. (**A**) Schematic maps of two sets of Cas9 constructs. The first four Cas9 constructs harbor a human-codon-optimized *SpCas9* (*Cas9*) (61) coding sequence (CDS) surrounded by two nuclear localization sequences (NLS-Cas9-NLS), linked to the *eGFP* CDS at its C-end via a self-cleaving T2A sequence, and terminated by the p10 3′-UTR from the *Autographa californica* nucleopolyhedrovirus (AcNPV) (48). The SpCas9 is expressed in early germ cells under *vasa* (vas) and *nanos* (*nos*) promoters, in late germ cells with *Bicaudal C* (*BicC*), and in both germ and somatic cells with *Ubiquitin 63E* (*ubiq)* promoter. These constructs also contain a red transgenesis marker (*Opie2-dsRed*). The second group of *Cas9* constructs carry a human-codon-optimized *SpCas9* expressed under the *vas* or *nos* promoter, and terminated with a single NLS (Cas9-NLS) and the corresponding *vas* and *nos* 3’UTR, as well as a green transgenesis marker (Ubiq-ZsGreen). (**B**) Images of homozygous transgenic SWD Cas9 ♀ flies generated with UbiqCas9.R and VasCas9.G. (**C**) Schematic maps of two gRNA constructs, and the targeted sequences in both *yellow* and *white* loci. The gRNA^y^ construct harbors the *yellow* gRNA (gRNA^y^) with a scaffold expressed with the *Dmel pU6-3* promoter and terminated by a PolyT terminator, and two green transgenesis markers, *Opie2-GFP* and *3xP3-GFP*. The gRNA^w^ construct harbors the *white* gRNA (gRNA^w^) with a scaffold expressed with the same *Dmel pU6-3* promoter and terminated the *pU6-3* terminator sequence, and a red transgenesis marker, *Ubiq-dsRed*. (**D**) Images of homozygous SWD *gRNA*^*y*^ and *gRNA*^*w*^ ♀. Both sets of RGB images for each ♀ fly were taken under the white light and corresponding fluorescent light illumination.

To genetically encode the gRNAs in SWD, we engineered two separate constructs encoding gRNAs targeting homozygous viable X-linked genes with recessive phenotypes: *yellow (y)* and *white* (*w*) (30, 31) (**Fig. 1C**). Each construct encoded one gRNA driven by the *D. melanogaster* small nuclear RNA *U6-3* promoter providing constitutive expression (49). The *gRNA*^*y*^ construct harbors two green transgenesis markers (*Opie2-eGFP* and *3xP3-eGFP*), and the *gRNA*^*w*^ construct contains one red marker (ubiq-dsRed). Both gRNA^y^ and gRNA^w^ constructs were assembled in *piggyBac* plasmids (**Fig. 1C**), and homozygous *gRNA*^*y*^ and *gRNA*^*w*^ lines were generated (**Fig. 1D, Table 1**).

### CRISPR-mediated mosaicism in F_1_ progeny

To assess the functionally of the strains produced, we genetically crossed homozygous *Cas9* ♀’s to homozygous *gRNA* ♂’s and examined expected eye and/or body coloration phenotypes in the resulting F_1_ progeny (**Fig. 2A-F; Table S1, S2**). This cross was performed to explore the rates of mutagenesis in the F_1_ somatic tissues, which is augmented by maternal deposition of Cas9 (45). High percentages (61.1%-100%) of F_1_ trans-heterozygous progeny generated by *Cas9*.*R* ♀’s crossed to *gRNA*^*y*^ ♂’s had visible yellow, instead of brown, body coloration indicating robust somatic *yellow* gene disruption (*y–* phenotype) (**Fig. 2B,C, Table S1**). Notably, when the *Cas9*.*R* ♀’s were crossed to *gRNA*^*w*^ ♂’s (**Fig. 2D**), we observed much higher variability (0% – 100%) for somatic *white* disruption in resulting F_1_ progeny (**Table S2**). While both sexes were affected, the majority of F_1_ trans-heterozygous progeny manifested highly variable mosaic eye coloration (*mW*), and a smaller fraction of F_1_ progeny had complete white eye color (*w–*, **Fig. 2E, F**). These results suggest that some somatic cells in mosaic flies harbored at least one wildtype *white* (*wt w+*) allele, or possibly a functional resistant (*w*^*R1*^) allele. Interestingly, the extensive somatic disruption of *yellow* and *white* was induced by all *Cas9*.*R* lines, while *Cas9*.*G* crossed to *gRNA*^*y*^ and *gRNA*^*w*^ strains did not appear to induce somatic mutations in the resulting F_1_ progeny (**Tables S1, S2**).

**Figure 2.**
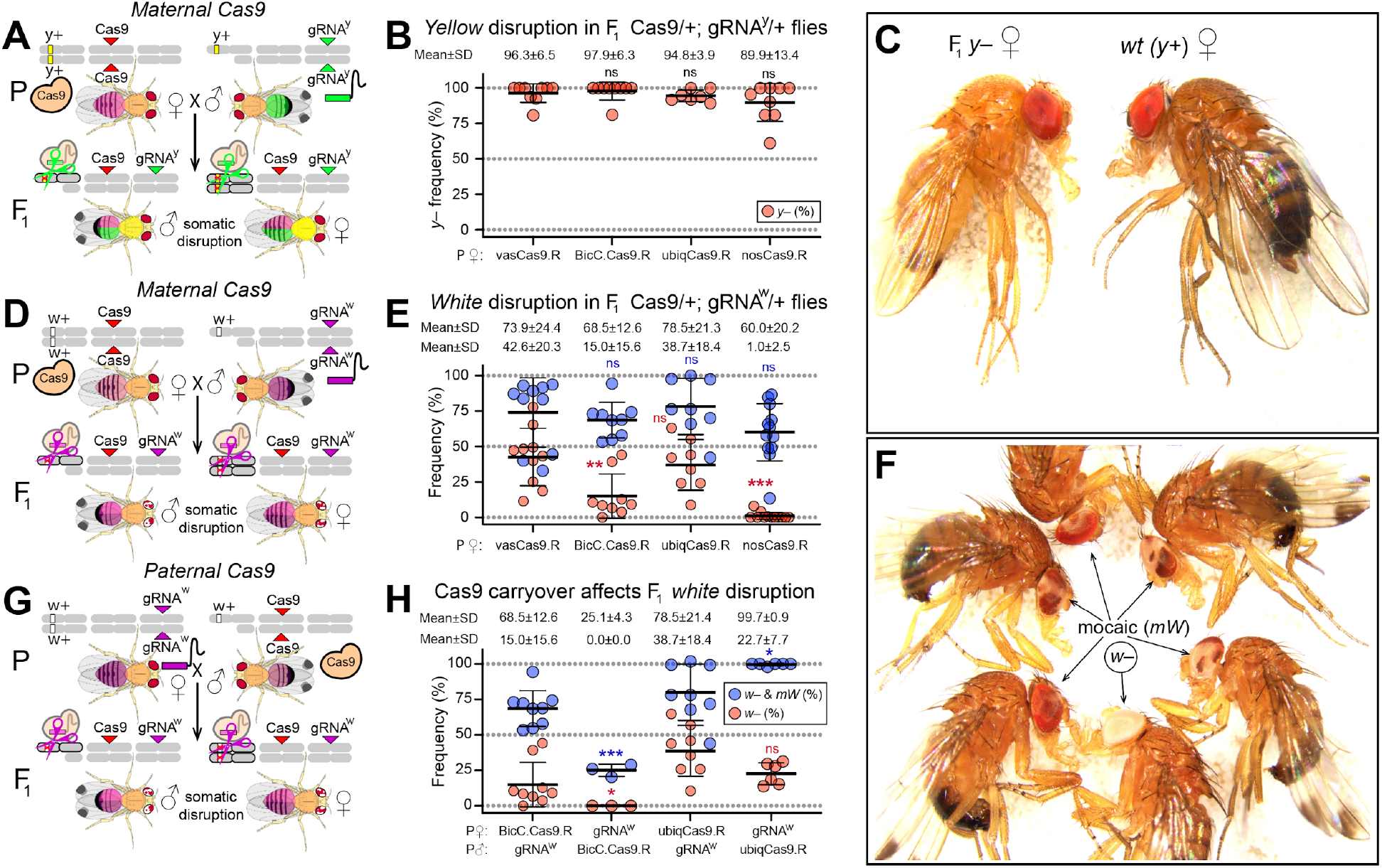
Disruption of *yellow* and *white* loci in F_1_ trans-heterozygous flies. (**A**) The schematic of a genetic cross between Cas9 and gRNA^y^ flies. To generate F_1_ trans-heterozygous flies, homozygous *Cas9* ♀ (red marker) crossed to homozygous *gRNA*^*y*^ ♂ (green marker). The targeted wildtype (*wt*) *yellow* gene (*y+* alleles, yellow stripes on X chromosome) is on the X chromosome. Yellow colored thoraxes in F_1_ flies indicate disruption of the *yellow* gene (*y-*). **(B)** Dot plot depicting the results of disruption of the *yellow* gene in somatic cells from F_1_ trans-heterozygous (*Cas9/+; gRNA*^*y*^*/+*) progeny using the Cas9.R strains inherited maternally. (**C**) Images of F_1_ *y–* trans-heterozygous and *wt y+* ♀. (**D**) The schematic of a genetic cross with maternal homozygous *Cas9* (red marker) and paternal homozygous *gRNA*^*w*^ (purple marker). The *white* gene is on the X chromosome (w+ alleles, white stripes on X chromosome). Red and white eye coloration in the F_1_ flies indicate somatic disruption of the *white* gene (*mW*). **(E)** Dot plot depicting the results of disruption of the *white* gene in somatic cells from F_1_ trans-heterozygous (*Cas9/+; gRNA*^*w*^*/+*) progeny using the Cas9.R strains inherited maternally. (**F**) Images of F_1_ trans-heterozygous mosaic *white* disruption (*mW*) and complete disruption (*w–*) phenotypes. (**G**) The schematic of a genetic cross between paternal Cas9 and maternal gRNA^w^ flies. To generate F_1_ trans-heterozygous flies, homozygous *Cas9* ♂ (red marker) crossed to homozygous *gRNA*^*w*^ ♀ (purple marker). The *white* gene is on the X chromosome (w+ alleles, white stripes). (**H**) Dot plot depicting the results of disruption of the *white* gene in somatic cells from F_1_ trans-heterozygous (*Cas9/+; gRNA*^*y*^*/+*) progeny using the Cas9.R strains inherited paternally. Plots show the mean ± SD over at least three biological replicates. Statistical significance was estimated using a two-sided Student’s *t* test with unequal variance. (*p* ≥ 0.05^ns^, *p* < 0.05*, *p* < 0.01**, and *p* < 0.001***).

### Maternal deposition and F_1_ Mosaicism

To determine whether maternal Cas9 deposition is essential to achieve high rates of F_1_ mutagenesis, we compared the rates of *white* mosaicism in F_1_ trans-heterozygous progeny inheriting Cas9 either maternally or paternally (**Fig. 2G,H**). We tested two Cas9 strains including *BicC-Cas9*.*R and ubiqCas9*.*R* which gave high rates of *white* mosaicism when Cas9 was provided maternally. The F_1_ *BicC-Cas9*.*R/+; gRNA*^*w*^*/+* with maternal *Cas9* showed significantly higher rates of *white* mosaicism then those with paternal *Cas9* (68.5±12.6% vs 25.1±4.3% of *w– & mW*, respectively; *p*<0.0002 *t-*test with equal variance, **Fig. 2H, Table S3**). Comparatively, the *ubiqCas9*.*R/+; gRNA*^*w*^*/+* with maternal *Cas9* showed significantly lower rates of *white* mosaicism then those with paternal *Cas9* (78.5±21.4% vs 99.7±0.9% of *w– & mW*, respectively, *p*<0.035 *t-*test with unequal variance, **Fig. 2H, Table S3**). Taken together, these data indicate that regardless of whether Cas9 was inherited maternally or paternally, high rates of mosaicism were observed in F_1_ trans-heterozygous progeny.

### Cas9/gRNA-mediated heritable mutations

To estimate the frequency of heritable germline mutations we mated F_1_ trans-heterozygous ♀’s harboring either maternal or paternal *Cas9/gRNA* to *wt* ♂’s, and assessed mutagenesis rates by scoring phenotypes in hemizygous F_2_ ♂’s progeny (**Fig 3A**). Both *yellow* and *white* are X-linked, therefore F_2_ ♂’s generated from F_1_ trans-heterozygous ♀’s inherit their mothers’ X chromosome. If the F_1_♀’s encode germline mutations in either *yellow* or *white*, then F_2_ ♂’s will inherit those X-linked mutations and display visible phenotypes (i.e. non-mosaic complete white eyes or yellow body). Of note, we specifically scored complete *yellow* and *white* phenotypes in F_2_ ♂ progeny here to differentiate heritable germline mutations from somatic mosaic mutations resulting from either maternal deposition or zygotic expression of transgenes. In combination with *gRNA*^*y*^, the *Cas9*.*R* strains induced significantly higher rates of *y–* alleles than those induced by the *Cas9*.*G* strains (77.6±12.5% and 2.0±2.1%, respectively; *p*<0.0001 *t-*test with unequal variance, **Fig. 3B,C, Table S1**). The F_1_ *ubiqCas9*.*R/+*; *gRNA*^*y*^*/+* ♀’s harboring maternal *Cas9* induced significantly higher levels of *y–* alleles than those carrying paternal *Cas9* (95.4±4.7% and 78.5±1.4, respectively; *p*<0.0003 *t-*test with equal variance), while the opposite was observed for F_1_ *vasCas9*.*G#6/+*; *gRNA*^*y*^*/+* ♀ (0.6±1.6% and 3.2±1.8, respectively; *p*<0.0163 *t-*test with equal variance, **Fig. 3B**). Interestingly, we observed higher variability of heritable *w–* mutations among replicates for the majority of *Cas9* strains (**Fig. 3D**). For example, independent groups of F_1_ *BicC-Cas9*.*R/+; gRNA*^*w*^*/+* or *vasCas9*.*G#4/+; gRNA*^*w*^*/+* ♀’s generated F_2_ ♂’s with LOF *w–* alleles in 92.7% and 1.1% of progenies, or those with *w–* in 83.5% and 0%, respectively (**Fig. 3D, Table S2**). Note that we did not find the mosaic eye coloration (*mW* phenotype, **Fig. 2F**) in F_2_ ♂, they had either *w–* or *w+* eye phenotype indicating the absence of somatic mosaicism (**Fig. 3E**).

**Figure 3.**
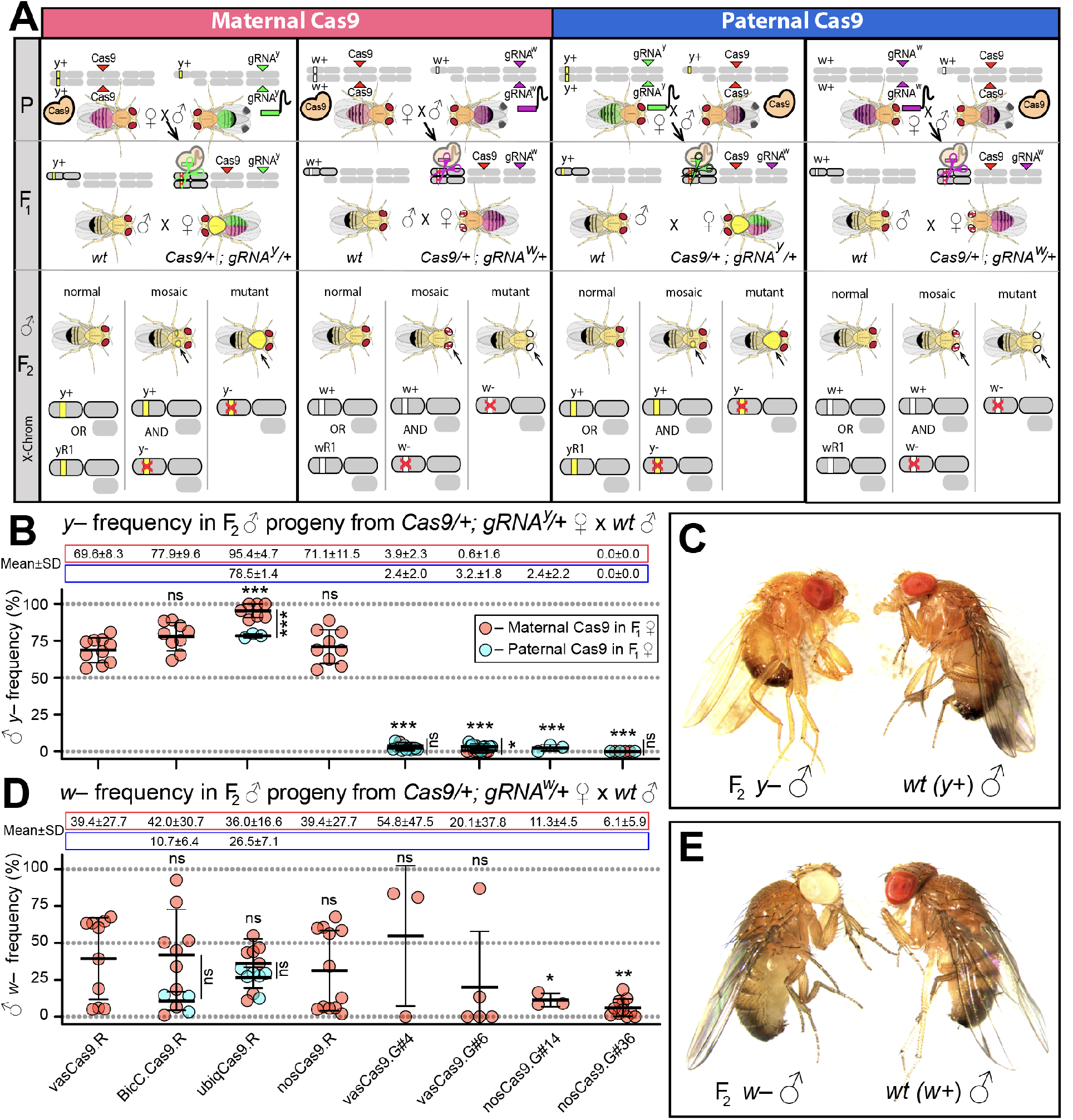
The efficiency of heritable *yellow* and *white* mutations. **(A)** To assess for heritable *yellow* or *white* mutations in germ cells, F_1_ trans-heterozygous *Cas9/+; gRNA/+* ♀’s were crossed to *wt* ♂’s, and *yellow* or *white* phenotype was scored in the generated F_2_ ♂ which inherited their mother’s X chromosome. Two types of trans-heterozygous ♀’s were used: the F_1_ ♀’s harboring maternal Cas9 (red points), and the F1 ♀’s harboring paternal Cas9 (blue points in B and D). (**B**) Dot plot depicting the *y–* frequency in the F_2_ ♂’s derived from crosses with the 8 homozygous Cas9 lines generated in this study. (**C**) Images of the generated F_2_ *y–* and *wt* (*y+*) ♂. (**D**) Dot plot depicting the *w–* frequency in the F_2_ ♂ derived from crosses with the 8 homozygous Cas9 lines generated in this study. (**E**) Images of the F_2_ *w–* and *wt* (*w+*) ♂. Plots show the mean ± SD over at least three biological replicates. Statistical significance was estimated using a two-sided Student’s *t* test with equal variance. (*p* ≥ 0.05^ns^, *p* < 0.05*, *p* < 0.01**, and *p* < 0.001***).

### Functional resistant alleles at the white locus

We observed higher variations in both F_1_ and F_2_ mutagenesis of *white* than compared to *yellow* (**Figs. 2B,E** and **3B**,**D**). We hypothesized that the F_2_ *w+* ♂ may harbor either uncut *wt w+* alleles or functional repaired alleles that were resistant to Cas9/gRNA^w^ cleavage (termed *w*^*R1*^ alleles) (**Fig. 4A**). To explore this hypothesis, we genotyped F_2_ *w+* ♂. Interestingly, we did not identify *wt w+* alleles among 37 F_2_ ♂ generated from multiple independent crosses. Instead, each genotyped F_2_ ♂ harbored either a one-base substitution (1bpSUB) or a twelve-base deletion (12bpΔ) directly at the *white* target sequence (**Fig. 4B**). Both in-frame *w*^*R1*^ alleles preserved *wt* function of the *white* gene. The A-to-G substitution in the 1bpSUB allele did not change the amino acid sequence (i.e. silent mutation), while the 12bpΔ allele contains a deletion of four amino acids (**Fig. 4B**). Notably, the 12bpΔ allele was sampled more frequently than the 1bpSUB allele, though both *w*^*R1*^ alleles were found in crosses with each *Cas9*.*R* strain. We also genotyped a few F_2_ *w–* ♂ and identified diverse LOF resistant alleles (termed *w*^*R2*^) induced by Cas9/gRNA^w^ (**Fig. 4C**). Each genotyped F_2_ *w–* ♂ harbored one LOF *w*^*R1*^ allele. Notably, we identified two in-frame LOF deletion alleles (3bpΔ and 12bpΔ*) that could not rescue the *wt white* function.

**Figure 4.**
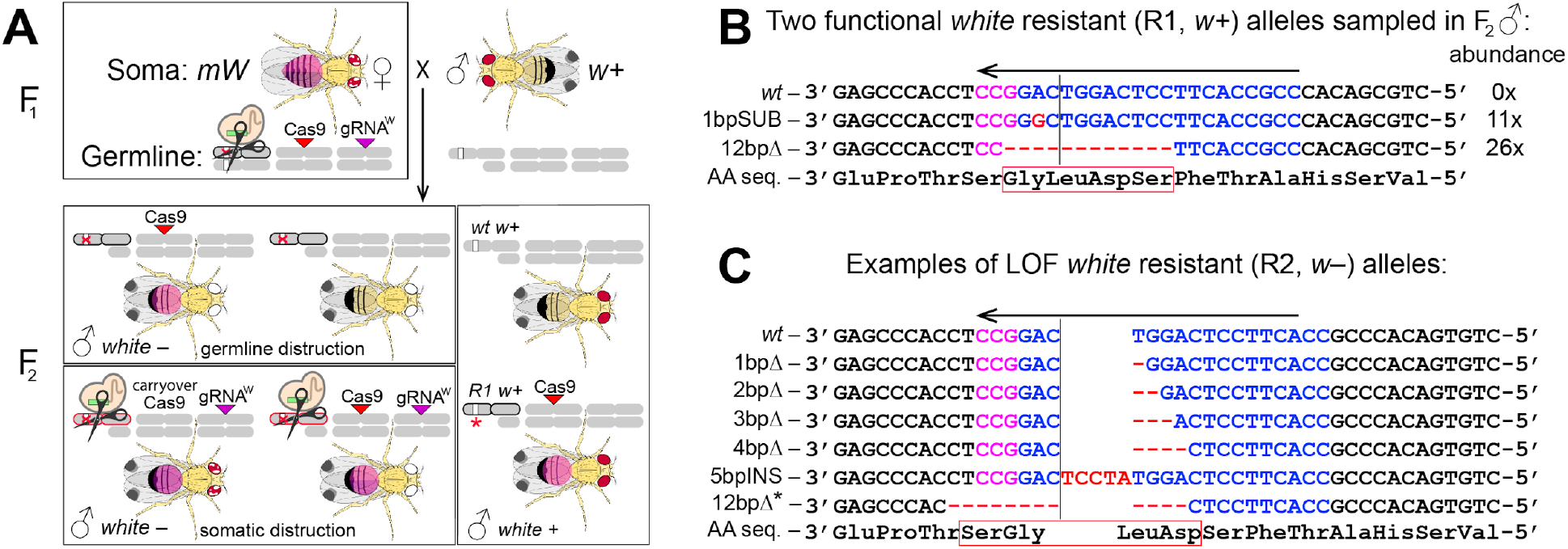
Functional and LOF resistant alleles at the *white* target loci. (**A**) Schematic depicting potential ♂ progeny from the genetic cross of F_1_ trans-heterozygous *Cas9/+; gRNA*^*w*^*/+* ♀ and *wt* (*w+*) ♂. The F_1_♀ has mosaic eye coloration (*mW*) indicating disruption of the *white* gene in its somatic tissues. Some *w+* alleles can be mutated in its germ cells resulting in F_2_ ♂ harboring the loss-of-function (LOF) *w–* allele (a crossed out white bar on the X chromosome with a black outline). Some *w+* alleles can be mutated in the somatic tissues of the F_2_ ♂ that inherited the *gRNA*^*w*^ transgene and have the carryover Cas9 protein (maternally deposited) or *Cas9* transgene (a crossed out white bar on the X chromosome with a red outline) also resulting in a LOF *w–* phenotype in F_2_ ♂. The F_2_ ♂ with normal eye coloration harbors the *wt w+* allele that escaped cleavage or the functional resistant alleles (R1, *w+*) that were mutated and became resistant to cutting by Cas9/gRNA^w^ but preserved the function of the *white* gene (a white bar with a red star on the X chromosome with a black outline). Note that LOF *w–* alleles contain insertions or deletions (*indels*) at the ligated cut site and are also likely resistant to Cas9/gRNA^w^, therefore they are referred to as R2 alleles. To explore the course of high variabilities in the *white* knockout, we sequenced the *white* target in F_2_ *w+* ♂. (**B**) Each genotyped ♂ from 37 F_2_ *w+* ♂ sampled from multiple independent crosses with each *Cas9*.*R* line had either one-base substitution (1spSUB) or twelve-base deletion (12bpΔ) directly at the *white* target sequence. The two R1 alleles persisted for two generations of cutting by Cas9/gRNA^w^, and maintained *wt* function of the *white* gene. No *wt w+* alleles were identified in any sampled F_2_ *w+* ♂. (**C**) LOS R2 allele sampled in F_2_ *w–* ♂. Note that the two sampled R2 alleles are in-frame (3bpΔ and 12bpΔ*), and yet they cannot restore the *wt* function. The sequence alignment of induced LOF alleles against the *wt* reference *white* sequence (58). The 20 bases of gRNA^w^ (in blue) and its PAM (purple) are depicted over the *white* target sequence. Arrows point the direction of the gRNA^w^ target. Mutated bases (red letters) and/or their absence (red dashes) are indicated relative to the *wt* sequence.

### Mutant y– strains were established

Both *yellow* and *white* genes have many features that makes them attractive targets and/or tools for genetic research. Therefore, we attempted to establish *w–* and *y–* strains to facilitate future genetic research in SWD. To this end, we generated independent F_2_ progeny (**Fig. 3**), and intercrossed individual F_2_ ♀ and ♂ harboring *y–* or *w–* mutant alleles in the absence of both *Cas9* and *gRNA* transgenes. We were able to establish and maintain eight homozygous *y–* strains, however we could not establish *w–* stocks as loss of function *w–* ♂ were sterile (30, 31). We genotyped each *y–* strain and identified five different insertions/deletions (*indel*) mutations at the *yellow* target sequence (**Fig. 5**). One six-base-deletion (6bpΔ) was induced independently in four *yellow* knockout strains (**Fig. 5**).

**Figure 5.**
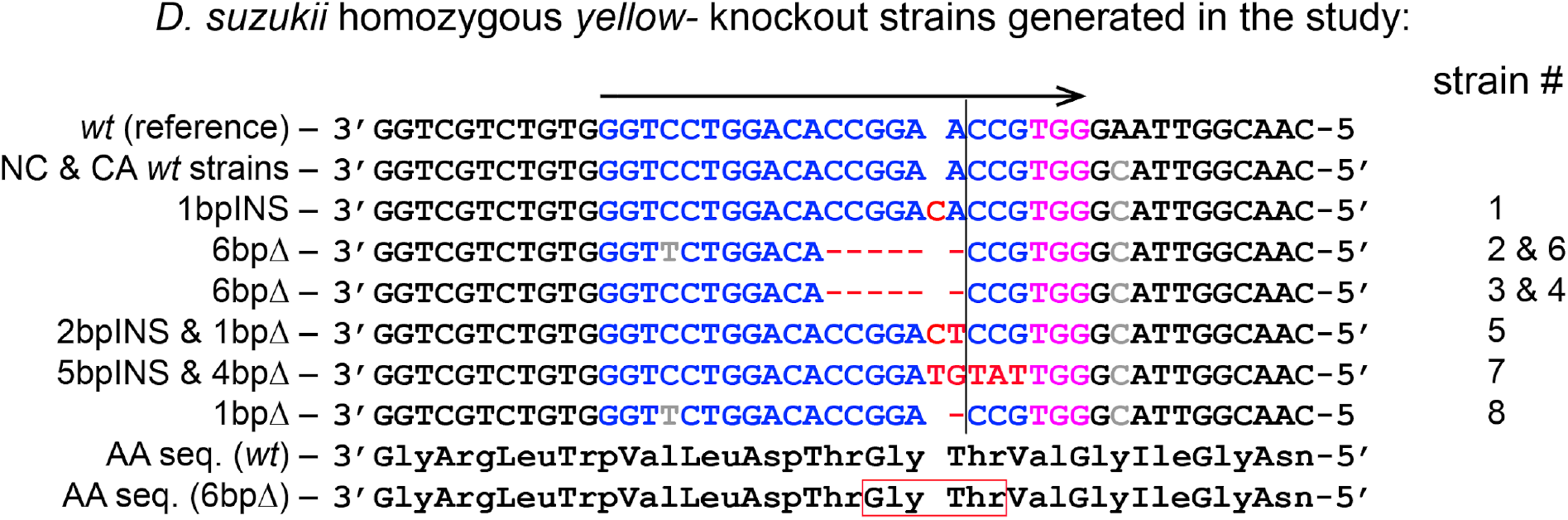
SWD *yellow* knockout (LOF) strains. We established eight independent homozygous viable *y–* knockout strains. The sequence alignment of induced LOF alleles against the *wt* reference *yellow* sequence (58). The 20 bases of gRNA are in blue, while the PAM sequence is in purple. The arrow points the direction of gRNA target. We identified two silent SNPs (grey colored base) in the SWD NC and CA *wt* stains relative to the *wt* reference strain (58): one at the 4th gRNA^y^ base and at the 2nd base after PAM silent the *yellow* targets. Mutated bases (red letters) and/or their absence (red dashes) are indicated relative to the *wt* sequence. The same six-base-deletion (6bpΔ) was independently induced in four strains #2, #3, #4, and #6. The 6bpΔ deletes two amino acids (Glycine and Threonine) that results in the LOF *y–* phenotype (**Figs. 2C, 3B**).

## Discussion

Here we describe the generation and assessment of an array of SWD transgenic strains expressing Cas9 driven under four promoters from *D. melanogaster*. To evaluate these Cas9 strains, two gRNA strains targeting X-linked recessive genes were also established. Using *Cas9* and *gRNA* strains, we demonstrate that the *Cas9*.*R* strains result in high rates of mutagenesis of targeted genes, while the *Cas9*.*G* strains had more limited efficacy.

Our results indicate that the *Cas9*.*R* constructs containing *NLS-Cas9-NLS* terminated with the p10 3’UTR resulted in the significantly stronger disruption of *yellow* in both somatic and germ cells than that by the *Cas9*.*G* harboring *Cas9-NLS* and terminated with the *vas* or *nos* 3’UTR. Due to random genomic integration of constructs, direct comparison of *Cas9* strains is confounded. Nevertheless, the observed differences were consistent among each *Cas9*.*R* and *Cas9*.*G* strain. The p10 3′-UTR from the *Autographa californica* nucleopolyhedrovirus (AcNPV) is known to increase efficiency of both polyadenylation and mRNA translation (47, 48). In addition, a seven-amino-acid-long SV40 NLS is shared between both *Cas9*.*R* and *Cas9*.*G* types, while *Cas9*.*R* constructs also contain a sixteen-amino-acid bipartite nucleoplasmin NLS (**Fig. 1A**). The inclusion of an additional NLS into Cas9 may provide increased nuclear localization. The *vas* and *nos* 3’UTRs used in the *Cas9*.*G* gene constructs would be expected to lead to localization of the Cas9 mRNA to the posterior end of the oocyte (50). In contrast, the p10 3’UTR is not predicted to result in such localization during oogenesis. Thus, our expectation was that the *Cas9*.*G* lines would be mostly active in germ cells that develop at the posterior end of the embryo, while the *Cas9*.*R* lines would have high activity in both somatic and germ cells. Several generated *Cas9*.*G* strains harboring independent insertions of *vasCas9*.*G* or *nos*.*Cas9*.*G* did not induce visible F_1_ somatic mutagenesis of *yellow* or *white* in the presence of the corresponding *gRNA*. Instead they caused the *yellow* or *white* mutagenesis in the F_2_ progeny indicating a possible germline-restricted Cas9 expression pattern.

We observed a high inter-replicate variability in frequencies of the *white* disruption in both somatic and germ cells. To explore cause of this variability, we genotyped many F_2_ *w+* ♂’s and found that each genotyped *w+* ♂ harbored one of the two functional *w*^*R1*^ alleles. Our results indicate that the Cas9/gRNA^w^ cleavage of *white* was efficient. We did not sample *wt w+* alleles in F_2_ ♂, instead they each harbored *w*^*R1*^ and *w*^*R2*^ resistant alleles. Both identified *w*^*R1*^ alleles, 1bpSUB and 12bpΔ, may have been induced by Cas9/gRNA^w^ or may have existed in the strains. The fact that the same two *w*^*R1*^ alleles were sampled in each cross with four *Cas9*.*R* strains also cannot rule out that at least one of them occurred naturally or was generated. However, the 12bpΔ allele, which removes 4 amino acids, potentially affects the fitness of its carriers, while the silent substitution is expected to be fitness neutral. The removal of 12 bases at the gRNA target site likely made the 12bpΔ allele much more resistant to the further Cas9/gRNA^w^-mediated cleavage than a single base substitution would; and the 12bpΔ allele was sampled more frequently than the 1bpSUB allele. Therefore, our data suggest that the 1bpSUB allele could pre-exist in SWD genetic background and might not be completely resistant to the Cas9/gRNA^w^, while the 12bpΔ allele was induced by Cas9/gRNA^w^ in our experiments.

Over a century ago, the establishment of *D. melanogaster* knockout strains of *white* and *yellow* provided valuable genetic background for seminal transgenic research (51). Previous studies demonstrated that unlike *D. melanogaster*, SWD ♂ harboring LOF *w-*alleles were sterile (30, 31), thus preventing the maintenance of the SWD *w–* strain. Here we describe the development of eight viable SWD *y–* strains. Five unique *y–* alleles were characterized from eight *y–* strains (**Fig. 5**). These strains may be useful reagents for applications in which fluorescent markers of transgenesis cannot be used, for example when genes of interest (GOI) themselves are linked to fluorescent tags. Therefore, the generated *y–* strains are valuable resources for genetic studies of SWD.

SWD is an invasive species and a close relative of the *D. melanogaster*, the classic genetic model organism for which a vast amount of detailed biological knowledge and a diverse array of genetic tools are available. Given the close phylogenetic proximity, the majority of know-how and genetic tools are easily portable across both *Drosophila* species and can be applied for the development of genetic methods for effective population control of SWD. The CRISPR/Cas9 technology has been used extensively for the precise genome editing in diverse animal and plant species. Both non-localized and localized methods for insect population control have been developed for *D. melanogaster* using the CRISPR/Cas9 technology (38, 40, 52, 53). The SWD tools strains described here should facilitate precise genetic engineering in the invasive pest species and may provide useful genetic reagents required for the development of genetic population control systems for SWD.

## Experimental Procedures

### Molecular Construct design and assembly

We used the previously described *piggyBac* plasmids harboring coding sequences of the *Cas9-T2A-eGFP* under different promoters and the *Opie2-dsRed* marker (38, 42, 45) to generate SWD *Cas9*.*R* strains: *vasCas9*.*R* (874Z plasmid, addgene #112687), *BicC*.*Cas9*.*R* (874R plasmid, addgene #168295), *UbiqCas9*.*R* (874W plasmid, addgene #112686), and *nosCas9*.*R* (874Z1 plasmid, addgene #112685). The Gibson enzymatic assembly method (54) or standard recombinant DNA methods were used to build the *piggyBac* transformation plasmids that carried *vasCas9*.*G* (addgene #169012), *nosCas9*.*G* (addgene #169011), *gRNA*^*y*^ (982G.2 plasmid, addgene #168294), and *gRNA*^*w*^ (addgene #169010) constructs. To assemble *vasCas9*.*G* and *nosCas9*.*G* driven by the *D. melanogaster vasa* and *nos* promoters, the vasa5’-Cas9-vasa3’ and nos5’-Cas9-nos3’ fragments were excised from *vasa-Cas9* plasmid DNA (Drosophila Genomics Resource Center #1340 (55)) and pBFv-nosp-Cas9 plasmid (32) (NIG-Fly, Japan), respectively, and then ligated into a *piggyBac* vector that was cut with HpaI and PspOMI. The *piggyBac* vector contains a ZsGreen fluorescent protein marker expressed by the *D. melanogaster polyubiquitin* gene promoter (*ubiq-ZsGreen*). This was made by excision of eGFP from MS1419 (56) and replacement with ZsGreen from pB[Lchsp83-ZsGreen] (57). We utilized previously described *yellow* (45) and *white* (31) gRNA sequences to build the *gRNA*^*y*^ and *gRNA*^*w*^ *piggyBac* plasmids (**Fig. 1C**). To assemble *gRNA*^*y*^, the *yellow* gRNA targeting an identical sequence in *D. melanogaster* ^(45)^ and SWD *yellow* exon#2 (DS10_00005318 (58)) was encoded into overlapping primers that amplified the *D. melanogaster U6-3* promoter on one side and the single chimeric gRNA scaffold (49, 59) with the poly(T)_6_ termination on the other side as previously described (38), and cloned into a piggyBac plasmid habording the *Opie2-eGFP-SV40* marker. Then, the additional *3xP3-eGFP-SV40* marker was added upstream from *U6-3-gRNA*^*y*^. To build *gRNA*^*w*^, the *U6:3-ex2* plasmid that contains *white* gRNA targeting SWD *white* exon#2 (DS10_00006062 (58)) and *D. melanogaster* U6:3 promoter and terminator (31) was digested with BglII and the excised U6:3p-gRNA-U6:3t fragment ligated with the *piggyBac* vector MS1425-p10 that contains a *ubiq-dsRed* marker. MS1425-p10 was derived from MS1425 (56) by excision of the SV40 pA sequence and replacement with p10 pA.

### SWD rearing and transgenesis

SWD were maintained on a cornmeal-yeast-agar diet at 21 °C with a 12/12h light/dark cycle at UCSD and/or NCSU. All test crosses were performed under these conditions except for those with the *vasCas9*.*G* and *nosCas9*.*G* lines and the *U6:3-ex2 white* gRNA lines that were performed at 25 °C. In both facilities, SWD flies were kept in an institutional biosafety committee-approved ACL1 (NCSU) and ACL2 (UCSD) insectaries and handled by limited expert investigators to prevent any unintended release of SWD. The SWD wildtype (*wt*) strains used in the study originated from Corvallis, OR (25) and North Carolina (31). Embryo injections were carried out at Rainbow Transgenic Flies, Inc. (http://www.rainbowgene.com) or at the NC State University insect transgenesis facility. Plasmids diluted in water to 200-300 µg/µl were injected into freshly collected embryos of the SWD harboring Hsp70Bb-piggyBacTransposase (26) or the North Carolina wild type strain with *piggyBac* helper plasmid(60). G_0_ adults emerging from injected embryos were outcrossed to SWD *wt* flies, and their G_1_ progeny were screened for the expression of specific fluorescent markers with the Leica M165FC fluorescent stereomicroscope.

### SWD genetics

To establish stable SWD transgenic lines that can be maintained over multiple generations, SWD expressing an independent insertion of a particular construct were repeatedly inter-crossed over multiple generations to generate a homozygous stock. The homozygosity of generated stocks was confirmed by test crosses using homozygous transgenic lines crossed to *wt* flies and scoring specific transgenic markers in their progeny. To our surprise, we observed that freshly eclosed SWD ♀ would frequently mate with older ♂; and, unlike *D. melanogaster*, the freshly eclosed phenotype by itself is not a sufficient indicator of the ♀ virginity. Therefore, to ensure that only virgin ♀ were used for genetic crosses, vials with eclosing flies were cleared up multiple times daily to remove old ♂.

We set the majority of genetic crosses in one direction, *Cas9* virgins ♀ were mated to *gRNA* ♂, to stimulate the gene disruption in somatic tissues of the F_1_ progeny by maternal Cas9 deposition (38) ^(45)^. Seven to ten homozygous Cas9 virgin ♀ were crossed to 7-10 homozygous gRNA ♂ in each vial, and the somatic disruption of *yellow* or *white* was scored in the F_1_ progeny (**Fig. 2A-F**). In addition to the complete absence of eye pigmentation (white eyes, *w–* phenotype), mosaic eye coloration was frequently observed in F_1_ trans-heterozygous *Cas9*.*R/+; gRNA*^*w*^*/+* flies of both sexes: it was recorded as a *mW* phenotype. We also performed genetic crosses with the paternal Cas9 for *BicC*.*Cas9*.*R* and *ubiqCas9*.*R* to assess the role of maternal Cas9 carryover on somatic gene disruption and to ascertain the insertion of *ubiqCas9*.*R* on the X chromosome, respectively (**Fig. 2G-H**). Both *yellow* and *white* genes are located on the X chromosome; and SWD ♂’s have only one X chromosome (i.e. hemizygous), which they inherit from their mothers. Therefore, to assess the mutation frequency in germ cells, 7-10 F_1_ trans-heterozygous virgin ♀ were crossed to 10 *wt* ♂, and *y–* or *w–* phenotype was scored in the F_2_ ♂ progeny. Flies were scored and imaged on the Leica M165FC fluorescent stereomicroscope equipped with the Leica DMC2900, View4K or Leica DFC500 camera.

### Genotyping the yellow and white target loci

To explore the molecular changes that caused resistant (R2) and functional in-frame resistant (R1) alleles. We PCR amplified a genomic region containing the target site for gRNA^y^ or gRNA^w^ (**Fig. 1C**) using single-fly genomic DNA preps (38)(45) from individual F_2_ ♂, which were scored for *yellow* or *white* phenotype. The 397bp PCR fragment of *yellow* target (exon #2) was amplified with 5’-GAATTCCAGCCACTCTGACTTATATCAATATGG-3’ (982G.s10F) and 5’-CAGGAGTAGGCAATTAAACCATAGCCC-3’ (982G.s11R); and 5’-GTGCCAGCACACGATCATCGGAGTGC-3’ (982A.s11R) and 5’-TGAGAAGAAGTCGACGGCTTCGCTGG-3’ (982A.s10F) primers were used to amplify the 322bp PCR fragment of *white* target (exon #2). PCR amplicons were purified using QIAquick PCR purification kit (QIAGEN), and sequenced in both directions with Sanger method at Genewiz, Inc. To characterize molecular changes at the targeted sites, sequence AB1 files were aligned against the corresponding reference sequences in SnapGene 4.

### Statistical analysis

Statistical analysis was performed in JMP8.0.2 by SAS Institute Inc. At least three biological replicates were used to generate statistical means for comparisons. P values were calculated for a two-sample Student’s t-test with equal or unequal variance. O’Brien’s test was used to assess that the variance is equal. All plots were constructed using Prism9 for macOS by GraphPad Software, LLC.

## Acknowledgments

We thank Amarish Yadav for molecular analysis of the *yellow* mutant lines. This work was supported in part by funding from Agragene Inc., California Cherry Board, and the Washington Tree Fruit Research Commission awarded to O.S.A. and from the National Institute of Food and Agriculture, U.S. Department of Agriculture Specialty Crops Research Initiative under agreement No. 2015-51181-24252 awarded to M.J.S. The views, opinions, and/or findings expressed are those of the authors and should not be interpreted as representing the official views or policies of the U.S. government.

## Ethical conduct of research

We have complied with all relevant ethical regulations for animal testing and research and conformed to the UCSD institutionally approved biological use authorization protocol (BUA #R2401).

## Author Contributions

O.S.A and M.J.S. conceived the study. N.P.K., E.J.B, F.L., A.Y., M.J.S. and A.B. designed experiments. N.P.K., EJ.B., A.Y. and A.B. obtained genetic cross data. A.B., T.Y., F.L. and I.S. designed and assembled constructs, A.B., F.L., and I.S. made transgenic lines, N.P.K. and J.L. performed molecular analyses. All authors approved the final manuscript.

## Data availability

Complete maps and Plasmid DNA assembled in the study were deposited at Addgene.org and available for distribution (**Table 1**). The SWD strains generated here will be made available upon request.

## Supporting Information

**Table S1**. *Yellow* disruption in F1 and F2 progeny (Cas9 x gRNA data).

**Table S2**. *White* disruption in F1 and F2 progeny (Cas9 x gRNA data).

**Table S3**. Cas9 protein carryover by *BicC*.*Cas9*.*R* ♀ of the *white* locus.

## References

1. D. B. Walsh, et al., Drosophila suzukii (Diptera: Drosophilidae): Invasive Pest of Ripening Soft Fruit Expanding its Geographic Range and Damage Potential. Journal of Integrated Pest Management 2, G1–G7 (2011).

2. A. Cini, C. Ioriatti, G. Anfora, A review of the invasion of Drosophila suzukii in Europe and a draft research agenda for integrated pest management. Bull. Insectology 54, 149–160 (2012).

3. O. Rota-Stabelli, M. Blaxter, G. Anfora, Drosophila suzukii. Curr. Biol. 23, R8–9 (2013).

4. A. B. Langille, E. M. Arteca, J. A. Newman, The impacts of climate change on the abundance and distribution of the Spotted Wing Drosophila (Drosophila suzukii) in the United States and Canada. PeerJ 5, e3192 (2017).

5. J. C. Lee, et al., Infestation of Wild and Ornamental Noncrop Fruits by Drosophila suzukii (Diptera: Drosophilidae). Annals of the Entomological Society of America 108, 117–129 (2015).

6. D. Mazzi, E. Bravin, M. Meraner, R. Finger, S. Kuske, Economic Impact of the Introduction and Establishment of Drosophila suzukii on Sweet Cherry Production in Switzerland. Insects 8 (2017).

7. D. A. Yeh, F. A. Drummond, M. I. Gómez, X. Fan, The Economic Impacts and Management of Spotted Wing Drosophila (Drosophila Suzukii): The Case of Wild Blueberries in Maine. J. Econ. Entomol. 113, 1262–1269 (2020).

8. M. P. Bolda, R. E. Goodhue, F. G. Zalom, Spotted Wing Drosophila: Potential Economic Impact of a Newly Established Pest. ARE Update 13, 5–8 (2010).

9. G. De Ros, S. Conci, T. Pantezzi, G. Savini, The economic impact of invasive pest Drosophila suzukii on berry production in the Province of Trento, Italy. Journal of Berry Research 5, 89–96 (2015).

10. L. A. Dos Santos, et al., Global potential distribution of Drosophila suzukii (Diptera, Drosophilidae). PLoS One 12, e0174318 (2017).

11. Season-long programs for control of Drosophila suzukii in southeastern U.S. blueberries. Crop Prot. 81, 76–84 (2016).

12. D. R. Haviland, E. H. Beers, Chemical Control Programs for Drosophila suzukii that Comply With International Limitations on Pesticide Residues for Exported Sweet Cherries. J Integr Pest Manag 3, F1–F6 (2012).

13. Control of spotted wing drosophila, Drosophila suzukii, by specific insecticides and by conventional and organic crop protection programs. Crop Prot. 54, 126–133 (2013).

14. F. R. M. Garcia, Drosophila suzukii Management (Springer Nature).

15. N. Desneux, A. Decourtye, J.-M. Delpuech, The sublethal effects of pesticides on beneficial arthropods. Annu. Rev. Entomol. 52, 81–106 (2007).

16. F. G. Z. Brian E Gress, Identification and risk assessment of spinosad resistance in a California population of Drosophila suzukii. Pest Management Science 75, 1270–1276 (2019).

17. E. F. Knipling, Possibilities of Insect Control or Eradication Through the Use of Sexually Sterile Males. J. Econ. Entomol. 48, 459–462 (1955).

18. A. P. Krüger, et al., Radiation effects on Drosophila suzukii (Diptera: Drosophilidae) reproductive behaviour. J. Appl. Entomol. 143, 88–94 (2019).

19. G. Lanouette, et al., The sterile insect technique for the management of the spotted wing drosophila, Drosophila suzukii: Establishing the optimum irradiation dose. PLoS One 12, e0180821 (2017).

20. H. Laven, A possible model for speciation by cytoplasmic isolation in the Culex pipiens complex. Bull. World Health Organ. 37, 263–266 (1967).

21. L. O’Connor, et al., Open release of male mosquitoes infected with a wolbachia biopesticide: field performance and infection containment. PLoS Negl. Trop. Dis. 6, e1797 (2012).

22. J. Engelstädter, A. Telschow, Cytoplasmic incompatibility and host population structure. Heredity 103, 196–207 (2009).

23. J. Cattel, et al., Back and forth Wolbachia transfers reveal efficient strains to control spotted wing drosophila populations. J. Appl. Ecol. 55, 2408–2418 (2018).

24. K. Nikolouli, F. Sassù, L. Mouton, C. Stauffer, K. Bourtzis, Combining sterile and incompatible insect techniques for the population suppression of. J. Pest Sci. 93, 647–661 (2020).

25. A. Buchman, J. M. Marshall, D. Ostrovski, T. Yang, O. S. Akbari, Synthetically engineered gene drive system in the worldwide crop pest. Proc. Natl. Acad. Sci. U. S. A. 115, 4725– 4730 (2018).

26. F.-C. Chu, W. Klobasa, N. Grubbs, M. D. Lorenzen, Development and use of a piggyBac-based jumpstarter system in Drosophila suzukii. Arch. Insect Biochem. Physiol. 97 (2018).

27. M. F. Schetelig, Y. Yan, Y. Zhao, A. M. Handler, Genomic targeting by recombinase-mediated cassette exchange in the spotted wing drosophila, Drosophila suzukii. Insect Mol. Biol. 28, 187–195 (2019).

28. H. M. M. Ahmed, F. Heese, E. A. Wimmer, Improvement on the genetic engineering of an invasive agricultural pest insect, the cherry vinegar fly, Drosophila suzukii. BMC Genet. 21, 139 (2020).

29. M. Paris, et al., Near-chromosome level genome assembly of the fruit pest Drosophila suzukii using long-read sequencing. Sci. Rep. 10, 11227 (2020).

30. P. Kalajdzic, M. F. Schetelig, CRISPR/Cas-mediated gene editing using purified protein in Drosophila suzukii. Entomol. Exp. Appl. 164, 350–362 (2017).

31. F. Li, M. J. Scott, CRISPR/Cas9-mediated mutagenesis of the white and Sex lethal loci in the invasive pest, Drosophila suzukii. Biochem. Biophys. Res. Commun. 469, 911–916 (2016).

32. S. Kondo, R. Ueda, Highly improved gene targeting by germline-specific Cas9 expression in Drosophila. Genetics 195, 715–721 (2013).

33. X. Ren, et al., Optimized gene editing technology for Drosophila melanogaster using germ line-specific Cas9. Proc. Natl. Acad. Sci. U. S. A. 110, 19012–19017 (2013).

34. S. J. Gratz, et al., Genome engineering of Drosophila with the CRISPR RNA-guided Cas9 nuclease. Genetics 194, 1029–1035 (2013).

35. Z. Yu, et al., Highly efficient genome modifications mediated by CRISPR/Cas9 in Drosophila. Genetics 195, 289–291 (2013).

36. O. Kanca, et al., An efficient CRISPR-based strategy to insert small and large fragments of DNA using short homology arms. Elife 8 (2019).

37. D. Li-Kroeger, et al., An expanded toolkit for gene tagging based on MiMIC and scarless CRISPR tagging in. Elife 7 (2018).

38. N. P. Kandul, et al., Transforming insect population control with precision guided sterile males with demonstration in flies. Nat. Commun. 10, 84 (2019).

39. M. Li, et al., Eliminating Mosquitoes with Precision Guided Sterile Males. bioRxiv, 2021.03.05.434167 (2021).

40. J. Champer, A. Buchman, O. S. Akbari, Cheating evolution: engineering gene drives to manipulate the fate of wild populations. Nat. Rev. Genet. 17, 146–159 (2016).

41. K. M. Esvelt, A. L. Smidler, F. Catteruccia, G. M. Church, Concerning RNA-guided gene drives for the alteration of wild populations. Elife 3 (2014).

42. M. Li, et al., Germline Cas9 expression yields highly efficient genome engineering in a major worldwide disease vector, Aedes aegypti. Proc. Natl. Acad. Sci. U. S. A. 114, E10540–E10549 (2017).

43. H. Sano, A. Nakamura, S. Kobayashi, Identification of a transcriptional regulatory region for germline-specific expression of vasa gene in Drosophila melanogaster. Mech. Dev. 112, 129–139 (2002).

44. M. Van Doren, A. L. Williamson, R. Lehmann, Regulation of zygotic gene expression in Drosophila primordial germ cells. Curr. Biol. 8, 243–246 (1998).

45. N. P. Kandul, et al., Assessment of a Split Homing Based Gene Drive for Efficient Knockout of Multiple Genes. G3 (2019) https://doi.org/10.1534/g3.119.400985.

46. O. S. Akbari, D. Oliver, K. Eyer, C.-Y. Pai, An Entry/Gateway cloning system for general expression of genes with molecular tags in Drosophila melanogaster. BMC Cell Biol. 10, 8 (2009).

47. M. M. van Oers, J. M. Vlak, H. O. Voorma, A. A. M. Thomas, Role of the 3’ untranslated region of baculovirus p10 mRNA in high-level expression of foreign genes. J. Gen. Virol. 80 (Pt 8), 2253–2262 (1999).

48. B. D. Pfeiffer, J. W. Truman, G. M. Rubin, Using translational enhancers to increase transgene expression in Drosophila. Proc. Natl. Acad. Sci. U. S. A. 109, 6626–6631 (2012).

49. F. Port, H.-M. Chen, T. Lee, S. L. Bullock, Optimized CRISPR/Cas tools for efficient germline and somatic genome engineering in Drosophila. Proc. Natl. Acad. Sci. U. S. A. 111, E2967–76 (2014).

50. R. Lehmann, C. Nüsslein-Volhard, The maternal gene nanos has a central role in posterior pattern formation of the Drosophila embryo. Development 112, 679–691 (1991).

51. A. H. Sturtevant, Experiments on sex recognition and the problem of sexual selection in Drosophila (1915) https://doi.org/10.1037/h0074109 (xMarch 11, 2021).

52. N. P. Kandul, J. Liu, J. B. Bennett, J. M. Marshall, O. S. Akbari, A confinable home and rescue gene drive for population modification. Elife 10 (2021).

53. G. Terradas, et al., Inherently confinable split-drive systems in Drosophila. Nat. Commun. 12, 1480 (2021).

54. D. G. Gibson, et al., Enzymatic assembly of DNA molecules up to several hundred kilobases. Nat. Methods 6, 343–345 (2009).

55. L. A. Baena-Lopez, C. Alexandre, A. Mitchell, L. Pasakarnis, J.-P. Vincent, Accelerated homologous recombination and subsequent genome modification in Drosophila. Development 140, 4818–4825 (2013).

56. M. F. Schetelig, X. Nirmala, A. M. Handler, Pro-apoptotic cell death genes, hid and reaper, from the tephritid pest species, Anastrepha suspensa. Apoptosis 16, 759–768 (2011).

57. C. Concha, et al., Efficient germ-line transformation of the economically important pest species Lucilia cuprina and Lucilia sericata (Diptera, Calliphoridae). Insect Biochem. Mol. Biol. 41, 70–75 (2011).

58. J. C. Chiu, et al., Genome of Drosophila suzukii, the spotted wing drosophila. G3 3, 2257– 2271 (2013).

59. A. R. Bassett, C. Tibbit, C. P. Ponting, J.-L. Liu, Highly Efficient Targeted Mutagenesis of Drosophila with the CRISPR/Cas9 System. Cell Rep. 6, 1178–1179 (2014).

60. X. Li, J. C. Heinrich, M. J. Scott, piggyBac-mediated transposition in Drosophila melanogaster: an evaluation of the use of constitutive promoters to control transposase gene expression. Insect Mol. Biol. 10, 447–455 (2001).

61. P. Mali, et al., RNA-guided human genome engineering via Cas9. Science 339, 823–826 (2013).

